# PRC1.6 localizes on chromatin with the human silencing hub (HUSH) complex for promoter-specific silencing

**DOI:** 10.1101/2024.07.12.603173

**Authors:** Tomás C. Rodríguez, Leonid Yurkovetskiy, Karthika Nagalekshmi, Chin Hung Oscar Lam, Eva Jazbec, Stacy A. Maitland, Scot A. Wolfe, Erik J. Sontheimer, Jeremy Luban

## Abstract

An obligate step in the life cycle of HIV-1 and other retroviruses is the establishment of the provirus in target cell chromosomes. Transcriptional regulation of proviruses is complex, and understanding the mechanisms underlying this regulation has ramifications for fundamental biology, human health, and gene therapy implementation. The three core components of the Human Silencing Hub (HUSH) complex, TASOR, MPHOSPH8 (MPP8), and PPHLN1 (Periphilin 1), were identified in forward genetic screens for host genes that repress provirus expression. Subsequent loss-of-function screens revealed accessory proteins that collaborate with the HUSH complex to silence proviruses in particular contexts. To identify proteins associated with a HUSH complex-repressed provirus in human cells, we developed a technique, Provirus Proximal Proteomics, based on proximity labeling with C-BERST (dCas9-APEX2 biotinylation at genomic elements by restricted spatial tagging). Our screen exploited a lentiviral reporter that is silenced by the HUSH complex in a manner that is independent of the integration site in chromatin. Our data reveal that proviruses silenced by the HUSH complex are associated with DNA repair, mRNA processing, and transcriptional silencing proteins, including L3MBTL2, a member of the non-canonical polycomb repressive complex 1.6 (PRC1.6). A forward genetic screen confirmed that PRC1.6 components L3MBTL2 and MGA contribute to HUSH complex-mediated silencing. PRC1.6 was then shown to silence HUSH-sensitive proviruses in a promoter-specific manner. Genome wide profiling showed striking colocalization of the PRC1.6 and HUSH complexes on chromatin, primarily at sites of active promoters. Finally, PRC1.6 binding at a subset of genes that are silenced by the HUSH complex was dependent on the core HUSH complex component MPP8. These studies offer new tools with great potential for studying the transcriptional regulation of proviruses and reveal crosstalk between the HUSH complex and PRC1.6.

## INTRODUCTION

The Human Silencing Hub (HUSH) complex is a key transcriptional regulator of endogenous retroelements and newly integrated proviruses in mammals [1,2]. Through direct interaction with target chromatin regions and nascent mRNA, HUSH complex core constituents, TASOR, MPHOSPH8 (MPP8), and PPHLN1 (Periphilin 1), selectively silence foreign and select endogenous genes at the pre- and post-transcriptional levels [3,4]. Additionally, HUSH recruits a variety of repressive complexes to genomic targets to establish durable, redundant silencing. In some cases, HUSH is associated with SETDB1, a histone methyltransferase that deposits trimethylated histone H3 at lysine 9 (H3K9me3) at constitutive heterochromatin [3,5], and MORC2, an ATPase that compacts chromatin [6]. A larger collection of accessory factors—including DNA methyltransferases [7], heterochromatin proteins [6,8], and nuclear export machinery [9]—may further explain the cell type specificity of HUSH silencing and its activity at some intron-containing endogenous genes [10].

Loss-of-function screens for human and mouse genes that promote silencing of retroviral, lentiviral, LINE-1, and heterologous transcriptional reporters, have repeatedly yielded the three core HUSH complex components, and conditionally, the HUSH complex accessory genes, SETDB1, ATF7IP, MORC2, and NP220 [3,6,8,11–14]. Additional genes required for HUSH complex-mediated silencing might elude forward genetic screens if they are essential or make only modest contributions to reporter repression. Proteomic approaches have offered alternative means to study the HUSH complex. Indeed, TASOR and MPP8 were identified as proteins associated with the primate immunodeficiency virus protein Vpx using stable-isotope labeling by amino acids in cell culture (SILAC)-based quantitative proteomics, coupled with liquid chromatography tandem mass spectrometry [15], as well as a targeted loss-of-function screen [14]. A yeast two-hybrid screen discovered that TASOR associates with the RNA deadenylase CCR4-NOT complex scaffold protein CNOT1 [16].

Spatially-restricted capture proteomics has been used to isolate proteins associated with the HUSH complex, independent of any effects on transgene expression. Specifically, TASOR was fused to an engineered biotin ligase (BirA*) to label and isolate proteins associated with the HUSH complex on chromatin [4]. In addition to the core HUSH complex components, this method identified NP220, as well as CCNK, MED19, RPRD2, FIP1L1, PPP1R10, and TOX4, proteins that regulate mRNA processing or RNA polymerase II activity [4]. A peptidomimetic compound targeting MPP8 was used to pull out MPP8, in complex with HP1, TRIM28, and HRP2 [17].

As a means to identify proteins that are specifically associated with a given genomic locus, we previously described a genomic locus-centered proximity proteomics tool, dCas9–APEX2 biotinylation at repetitive genomic elements by restricted spatial tagging (C-BERST) [18]. In this method, the ascorbate peroxidase APEX2 is recruited to a genomic locus of interest by fusing it to Cas9 in which the two nuclease active sites are disrupted by mutation (dCas9). In the presence of biotin-phenol and H_2_O_2_, APEX2 generates biotin-phenoxy radicals that attach covalently to tyrosine residues within a radius of ∼20 nm. We used this tool to map the proteome around centromeres and telomeres [18]. Recently, this proximity labeling strategy was used to characterize the local protein environment of evolutionarily young LINE1 elements: among the core HUSH complex components, only PPHLN1 was identified in this screen [19]. These results demonstrate the feasibility of using proteomics to identify proteins at HUSH complex-silenced loci, independent of each factor’s contribution to HUSH complex function.

Here, we developed a modified version of C-BERST, Provirus Proximity Proteomics, to assess the proteins near a provirus. The screen was carried out in the presence and absence of the core HUSH component, MPP8, using spatially restricted protein tagging of proteins in close proximity to an efficiently silenced, HUSH complex-sensitive reporter. A number of host proteins were found to be associated with the HUSH complex-silenced proviruses, including DNA repair, mRNA processing, and transcriptional silencing proteins. We describe the complex balance between transcription-promoting and repressing factors recruited in the presence of the HUSH complex. We show that the localization of L3MBTL2, a component of the non-canonical polycomb repressive complex 1.6 (PRC1.6), is highly correlated with that of HUSH complex components, both at specific reporters and genome-wide. Additionally, using an orthologous, genetic loss-of-function screen with a HUSH-sensitive reporter, we identified L3MBTL2 and the PRC1.6 gene MGA. PRC1.6 was found to participate in silencing at a subset of HUSH complex-sensitive lentivectors. Finally, the proteomics and global chromatin mapping data were integrated to better characterize the unique chromatin environment common to PRC1.6 and HUSH complex-bound loci.

## RESULTS

### Site-specific proteomics of a HUSH complex-silenced provirus

The expression of transgenes delivered to cells by lentiviral, retroviral, or AAV vectors is often silenced [3,8,20]. While such transcriptional repression depends on multiple factors, including the host cell type, the location within the chromatin of the integration site, the length and deoxynucleotide content of the transgene, and the presence or absence of *cis*-acting splice signals, transcriptional silencing of these transgenes is often caused by the HUSH complex and its cofactors [1,2,10]. We developed transducible transcriptional reporters that are silenced by the HUSH complex in a manner that is independent of the site of integration in the chromatin, and leveraged this system to identify host genes involved in HUSH complex function [14,21]. Position-independence of these HUSH-sensitive reporters has the advantage that cell sorting is not required to isolate the silenced clones from a heterogeneous pool of transductants, as was the case in previous screens [3]. This technical detail was important to the experiments described here. Our previous experiments indicated that targeting of APEX2 to a chromosomal locus by fusing it to dCas9 worked well to map the proteome of centromeres and telomeres - which are present at multiple copies per cell - but lacked the sensitivity to characterize the protein-environment of single copy genes [18]. We reasoned that, given the position-independence for HUSH complex-mediated silencing of our reporters, proviruses at diverse loci would be silenced by the HUSH complex via a similar mechanism. Therefore, these reporters could be used to generate a pool of clones at high multiplicity of infection to boost the sensitivity of detection by dCas9-directed APEX2.

To test this hypothesis, we profiled the proteins enriched at a pool of proviruses generated by transduction with a given reporter (Provirus Proximity Proteomics), in the presence or absence of the core HUSH complex protein, MPP8. First, HEK293T cells were transduced with a previously described lentiviral vector bearing doxycycline-inducible dCas9-mCherry-APEX2 fused to an FKBP degron (Figure 1A) [18]. An sgRNA targeting a sequence common to the 5’ and 3’ LTRs of the reporter (sgLTR) was introduced next using the *Sleeping Beauty* transposon [22]. As a control, to subtract the background of nuclear proteins, cells were generated in parallel with an sgRNA (sgNS) that does not target sequences in the human genome and has been shown to distribute uniformly throughout the nucleus [18]. Cells were then transduced with a lentiviral vector that expresses an *MPP8*-specific shRNA to knockdown MPP8 mRNA and abrogate HUSH complex function; these cells are indicated as HUSH^-^ in Figure 1A. A control population was generated by transduction with an shRNA that targets luciferase (shLuc [23]), indicated as HUSH^+^ in Figure 1A. Next, cells were transduced at a multiplicity of infection of five with a lentivector, pscALP-PLXIN-*gag*GFP-WPRE, that drives a previously-described *gag*-GFP fusion gene [14] from an internal MLV LTR. This latter vector exhibits integration site-independent, HUSH complex-sensitive, transcriptional repression in HEK293T cells, as assessed for GFP mean fluorescence intensity by flow cytometry and by RT-qPCR for reporter RNA (Supplementary Figure 1).

**Figure 1:**
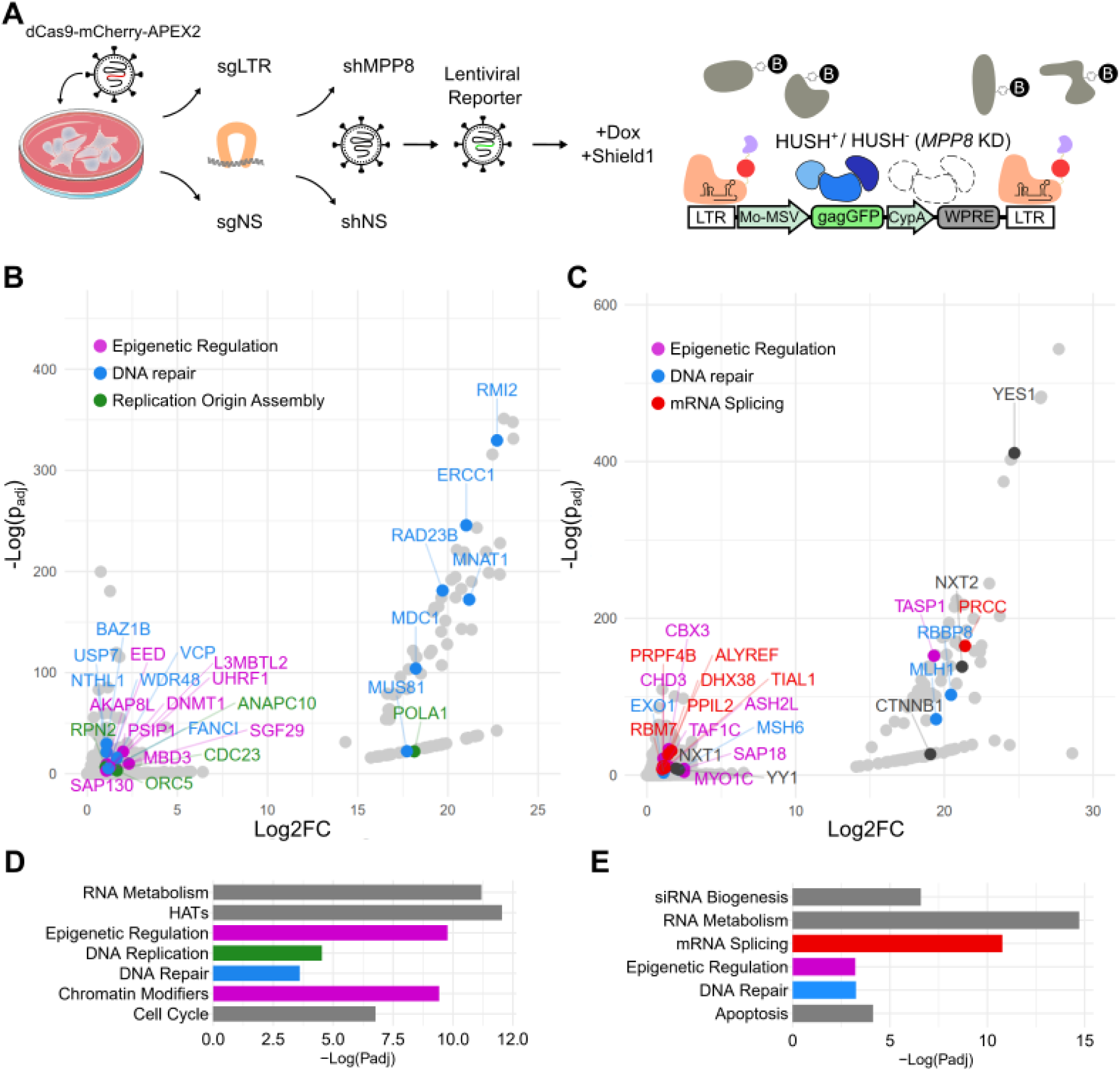
Provirus Proximity Proteomics in human cells. (A) Schematic describing the Provirus Proximity Proteomics approach targeting the provirus of a lentivector after transduction of HEK293T cells. An sgRNA targeting the provirus LTRs (sgLTR) recruited dCas9-mCherry-APEX2, then a pulse of biotin-phenol was conjugated to proteins in the immediate vicinity. HUSH^-^ indicates cells transduced with an shRNA targeting MPP8 (*MPP8* KD). HUSH^+^ indicates cells transduced with a non-targeting control shRNA. (B,C) Proteins enriched at the provirus (sgLTR) in HUSH^+^ and HUSH^-^ cells, respectively. Named proteins are significantly enriched over non-specific nuclear background (p_adj_ < 0.05; Log2FC > 1) and colored according to the function annotated in D and E. (D,E) Enrichment of Reactome [24] pathways in HUSH^+^ and HUSH^-^ backgrounds, respectively. Colors correspond to member proteins in panels B and C. For information associated with this figure please see Extended Data Tables 1, 2, 3, 4, and 5.

Once the multiply-transduced cell lines were established, transcription of dCas9-mCherry-APEX2 was induced for 24 hrs with doxycycline, and the APEX2-degron fusion protein was stabilized for the last 16 hrs with Shield1 ligand, as previously described [18]. Cells were incubated with biotin-phenol and APEX2-mediated proximity labeling was activated with H_2_O_2_ treatment for 1 minute. The reaction was quenched with sodium ascorbate and cells were washed with PBS. Nuclei were isolated and sonicated, and biotinylated proteins were enriched on streptavidin dynabeads. Proteins were eluted using SDS and identified by mass spectrometry.

Peptides from the three core HUSH components, MPP8, TASOR, and PPHLN1, were detected in both the shLuc control knockdown cells (HUSH^+^ in Figure 1A) and in the *MPP8* knockdown cells (HUSH^-^ in Figure 1A), but the abundance of these peptides was much greater in the HUSH^+^ condition (Supplementary Figure 2; Extended Data Table 1). As compared with the sgRNA nuclear background signal (sgNS), HUSH^+^ and HUSH^-^ cells bearing sgLTR showed significant enrichment of 229 and 259 uniquely identifiable peptides, respectively (Figure 1B and C; p_adj_<0.05, n=3 replicates per condition; Extended Data Tables 2 and 3). The majority of the significantly enriched proteins localize to the nucleus, indicating that these data are unlikely to represent contamination from off-target cellular compartments. In aggregate, proteins from both HUSH^+^ and HUSH^-^ conditions overrepresent several reactome pathways corresponding to nucleus-specific functions, including RNA metabolism, epigenetic regulation, DNA repair (Figure 1D and E; Extended Data Tables 4 and 5).

Despite efficient silencing in HUSH^+^ cells, mRNA processing and transcription-related pathways were among the most significantly enriched (Figure 1B and Extended Data Table 2). For example, NuA4-related histone acetyltransferase complex proteins YEATS4, MEAF6, YEATS4, and MORF4L, as well as MSL complex proteins MSL1 and MSL3, were identified in these cells. General transcription machinery such as MNAT1, TAF12, SUPT5H, and SGF29 were also enriched. Transcriptionally repressive pathways were also found as hits in HUSH^+^ cells, including the DNA methylation-associated factors RB1 and DNMT1 and nuclear export-related factors NUP85 and SARNP, HIV-1 integration regulators CTNNBL1 and PSIP1, polycomb repressive complex factors including EED, PCGF5, and L3MBTL2, and the histone deacetylation factors SAP130, ING2, MBD3, and ANP32E. Importantly, we also found UHRF1, which binds H3K9me2/3 directly and recruits both histone deacetylases, histone methyltransferases (G9a), and DNA methyltransferases (DNMT1) [25].

A constellation of transcriptionally activating and repressing complexes in the provirus-associated proteome of HUSH^−^ cells was observed despite much greater reporter expression in these cells than in the control HUSH^+^ cells (Figure 1C and Extended Data Table 3). Interestingly, SETDB1 and CBX3 heterochromatin proteins were among the enriched factors. SETDB1, an H3K9 methyltransferase, is necessary for HUSH-mediated silencing of diverse endogenous loci and reporters [3]. CBX3 binds methylated H3K9, is found primarily at constitutive heterochromatin, suppress MMTV expression [26], and maintains HIV-1 latency [27], though it has been associated with genic regions and post-transcriptional RNA processing [28]. YY1 is also enriched and may explain the recruitment of SETDB1 as part of the HUSH-independent TRIM28/KAP1 silencing pathway [29]. We also identified NXT1, NXT2, and ALYREF, all of which participate in the nuclear export of spliced mRNAs and are also necessary for export of retroviral transcripts and replication [30,31]. Compared to HUSH^+^ cells, several spliceosome-associated factors, associated with major and minor splicing pathways, were enriched in HUSH^-^ cells (Figures 1B and C; Extended Data Tables 2 and 3). This was particularly interesting given recent evidence that HUSH and the spliceosome may have antagonistic activities [10]. Several KRAB zinc finger proteins (KZNFs), ZBTB17, ZBTB7A, ZNF184, ZNF280C, ZNF281, ZNF451, ZNF668, ZNF687, ZNF691, and ZNF845, were enriched at the provirus in HUSH^-^ cells. As KZNF genes are known endogenous HUSH targets [6], we postulate that KZNFs are de-repressed after *MPP8* knockout and bind provirus as part of the human innate defense to retroviruses and parasitic endogenous retroelements [32]. Finally, we note that two potent transcriptional activators, YAP1 and CTNNB1, are found in the HUSH^-^ condition and likely drive some of the remodeling and transcriptional activity that was observed (Supplementary Figure 1).

### Unique DNA repair, replication, and polycomb repressive factors define the HUSH-associated proteome

Peptides from 179 proteins were exclusively enriched at the provirus in the HUSH^+^ condition. To better understand which of these proteins contribute to transcriptional silencing at endogenous loci, these hits were compared to those from published proximity labeling datasets. Previously, C-BERST was used to target and profile evolutionarily “young” LINE1 elements in two cell types, LNCaP and E006AA-hT [19]. Considering that many transcriptionally active LINE1s are HUSH targets [6,13], we reasoned that proteins enriched at both young LINE1s and provirus may interact with HUSH to carry out core repressive functions (Figure 2A). Among overlapping factors, we noted recurrent pathways including DNA repair (MDC1, ERCC1, and SMC5), chromatin remodeling (MEAF6 and VPS72), DNA methylation (DNMT1), and DNA replication (CDC23). We next compared HUSH^+^ hits to factors enriched in U2OS using APEX2-MDC1, finding 10 overlapping proteins including ERCC1 and SMC5. As HUSH and MDC1 were previously shown to resolve breaks in ribosomal DNA [33], we posit that these factors are important for silencing or repair of HUSH-sensitive loci genome-wide. Finally, we compared protein hits from HUSH^+^ cells to those found at constitutive heterochromatin by H3K9me3 ChromoID [34]. BAZ1B, DNMT1, PSIP1, and UHRF1 were shared in both proteomes, suggesting that HUSH may recruit these factors to establish robust silencing at target loci.

**Figure 2:**
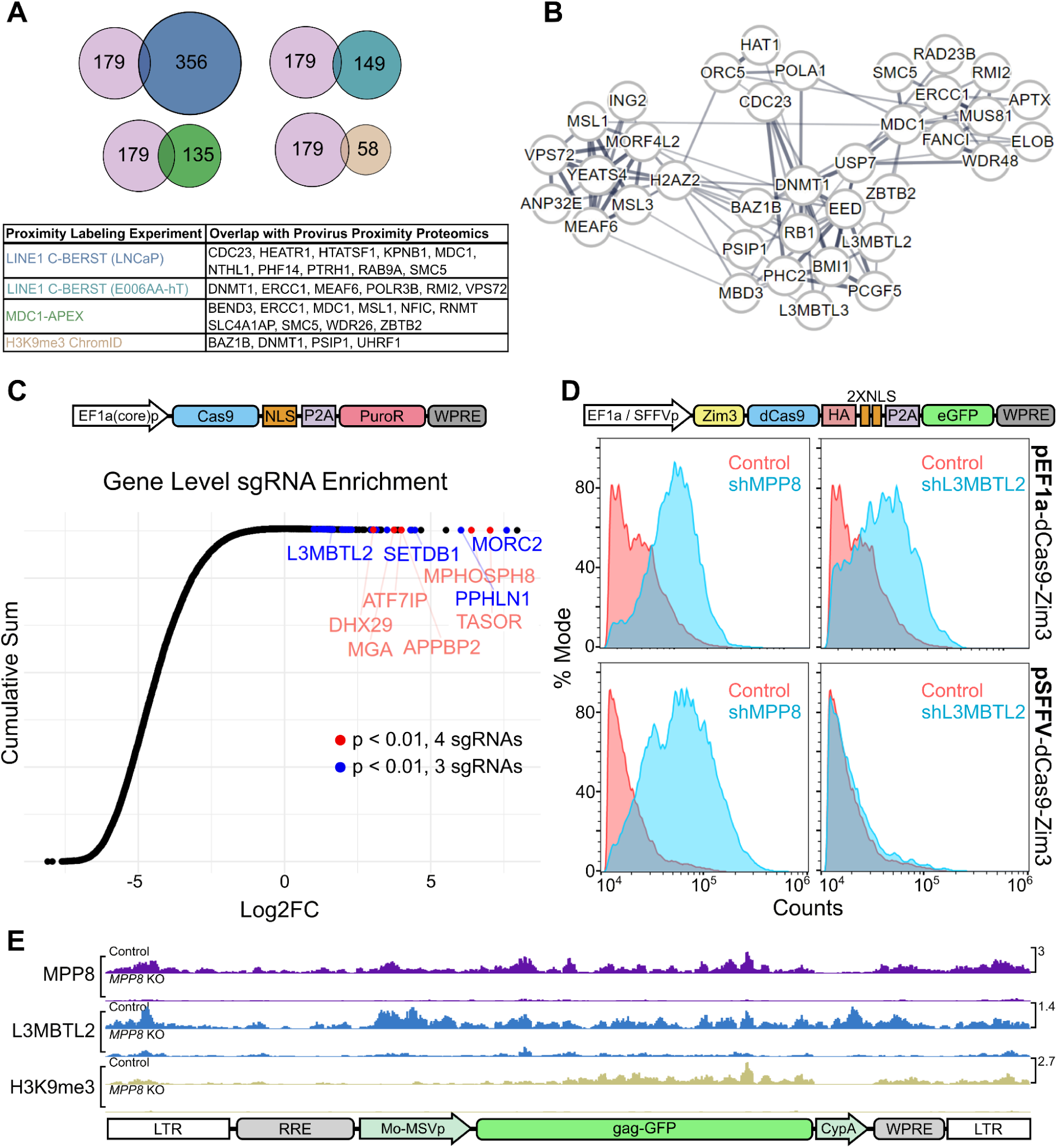
Validation of HUSH^+^ proteins enriched by Provirus Proximity Proteomics using orthogonal assays. (A) Comparison of the 179 proteins enriched at the provirus under HUSH^+^ conditions with those proteins enriched in published proximity labeling datasets [19,34,36]. (B) STRING network analysis of proteins enriched at the provirus in HUSH^+^ cells. Line widths are in proportion to the evidence supporting interaction between two nodes, as described in [35]. (C) CRISPR knockout screen conducted in Jurkat cells using single vector Toronto KnockOut (TKO) CRISPR library Version 3 with a puromycin selection cassette. Cumulative sum plot depicts the comparison of sgRNA relative quantity in cell populations transduced with library and treated with 10 μg puromycin versus a population transduced with library and treated with 1 μg puromycin. Individual Log2FC values represent aggregate counts of multiple sgRNAs corresponding to a single gene calculated by MAGeCK [37]. Red and blue points meet significance criteria (p < 0.01) for 3 (blue) or 4 (red) sgRNAs. (D) Flow cytometry histogram depicting GFP-positive population of Jurkat cells transduced with pEF1a-Zim3-dCas9-P2A-EGFP (top panels) or pSFFV-Zim3-dCas9-P2A-EGFP (bottom panels). Each population was treated with shRNA vectors targeting *MPP8* (left panels) or *L3MBTL2* (right panels) in blue, vs the non-targeting shRNA control in red. (E) Normalized CUT&Tag signal (counts-per-million) for MPP8, L3MBTL2, and H3K9me3 at the pscALP-PLXIN-gagGFP-WPRE provirus in HEK293T cells. For data associated with this figure please see Extended Data Table 6 and SRA Bioproject PRJNA869850.

Using STRING [35], we searched for HUSH^+^-enriched proteins related to those found in the young LINE1 element proteomes (Figure 2B). We observed networks of highly interconnected nodes associated with MDC1/ERCC1, DNMT1, VPS72/MEAF6, and CDC23. Polycomb group proteins EED, PCGF5, PHC2, and L3MBTL2 were located at the center of these networks despite evidence that HUSH-silenced loci do not accumulate H3K27me3 deposited by canonical polycomb complexes [3]. These intriguing findings led us to examine the role of polycomb group proteins in the repression of HUSH-sensitive DNA elements.

### PRC1.6 is necessary for silencing of a HUSH-sensitive transgene expressed from the pEF1a promoter

Forward genetic screens for host factors necessary for silencing of newly-integrated retroviruses have implicated the HUSH complex core components (MPP8, TASOR, and PPHLN1) and the HUSH accessory proteins (SETDB1, ATF7IP, and MORC2, among others) [3,6,8,10]. Enrichment of these factors was dependent on reporter/transgene sequence content, provirus location, and cell type. We identified a lentiviral reporter gene with a long open-reading frame that is HUSH-dependent for silencing in a vast pool of clones, indicating that its HUSH-sensitivity was independent of the site of integration [14]. Similarly, we found that the large transgene Cas9 is efficiently silenced by the HUSH complex, as others have also found [10]. We thus conducted a CRISPR loss-of-function screen using a construct in which Cas9 expression is coupled with puromycin resistance via the P2A peptide. By adding 10x higher puromycin, as compared to the standard levels of puromycin used for selection, we reasoned that sgRNA guides enhancing Cas9-P2A-Puro expression would also allow for selection based on loss of HUSH silencing. In Jurkat cells, this screen identified sgRNAs targeting genes encoding core HUSH complex components (TASOR, MPP8, and PPHLN1), the previously characterized HUSH accessory proteins (SETDB1 and ATF7IP), and the PRC1.6 components L3MBTL2 and MGA (Figure 2C and Extended Data Table 6).

Next, we asked whether Cas9-containing transgenes are silenced by PRC1.6. To address this question, we transduced Jurkat cells with a lentivector harboring pEF1a-Zim3-dCas9-P2A-GFP [38]. We then monitored GFP expression after transduction with *MPP8*- and *L3MBTL2*-targeting shRNAs. Compared to cells transduced with a non-specific shRNA, both *MPP8* and *L3MBTL2* knockdowns demonstrated a similar increase in GFP expression (Figure 2D).

HUSH silencing of lentivectors is modulated at least in part by the promoter that drives expression of the transgene [3,8,14,21]. Both the Cas9-P2A-Puro vector used in our loss-of-function screen (Figure 2C), and the Zim3-dCas9 vector that was used to confirm the effect of L3MBTL2 (Figure 2D), are driven by EF1a promoters. The core EF1a promoter is used in the former, and the full EF1a promoter with intron 1 is used in the later. To understand if HUSH- and PRC1.6-mediated repression is specific to EF1a, we transduced Jurkats with pSFFV-Zim3-dCas9-P2A-GFP, an isogenic vector that has only the EF1a promoter replaced with the SFFV promoter [38]. After transduction with shRNA vectors, we observed no GFP signal change in the *L3MBTL2* knockdown whereas the *MPP8* knockdown continued to demonstrate increased GFP compared to control (Figure 2D), indicating that the pSFFV-driven transgene is not repressed by PRC1.6.

### L3MBTL2 localization to the lentiviral provirus is MPP8-dependent

After demonstrating that PRC1.6 represses HUSH-sensitive transgenes under pEF1a control, we sought to understand whether PRC1.6 components localize directly to the provirus that was used in our Provirus Proximal Proteomics screen (Figure 1). HEK293T cells were first transduced with pscALP-PLXIN-gagGFP-WPRE. Then, MPP8 was depleted by CRISPR knockout. Consistent with the results of the Provirus Proximity Proteomics screen, Cleavage Under Targets and Tagmentation (CUT&Tag) demonstrated that MPP8 and L3MBTL2 are enriched at the provirus in control cells (Figure 2E and SRA Bioproject PRJNA869850). In contrast, the signal for both factors was substantially depleted in cells with MPP8 knocked out. Additionally, the marker for transcriptional repression, H3K9me3, was also abrogated in the absence of MPP8.

### PRC1.6 co-localizes with HUSH at endogenous loci

While PRC1.6 and HUSH co-localize to and silence select transgenes, it is unclear whether they share similar endogenous targets. To explore this, we generated Cas9-induced *L3MBTL2* knockouts in Jurkat cells and compared transcriptome-wide gene expression with that of cells treated with a non-specific sgRNA (Figure 3A and Extended Data Table 7). Among significantly (p_adj_ < 0.1) upregulated transcripts, we noted *EEF1AP10*, an intronless EF1a pseudogene. *TASOR*, KRAB zinc fingers, and intronless genes are also upregulated in *L3MBTL2* knockout cells. Surprisingly, none of these transcripts are significantly overexpressed in *MPP8* knockout Jurkats, suggesting that PRC1.6 and HUSH do not collaborate to silence endogenous genes in this cell type.

**Figure 3:**
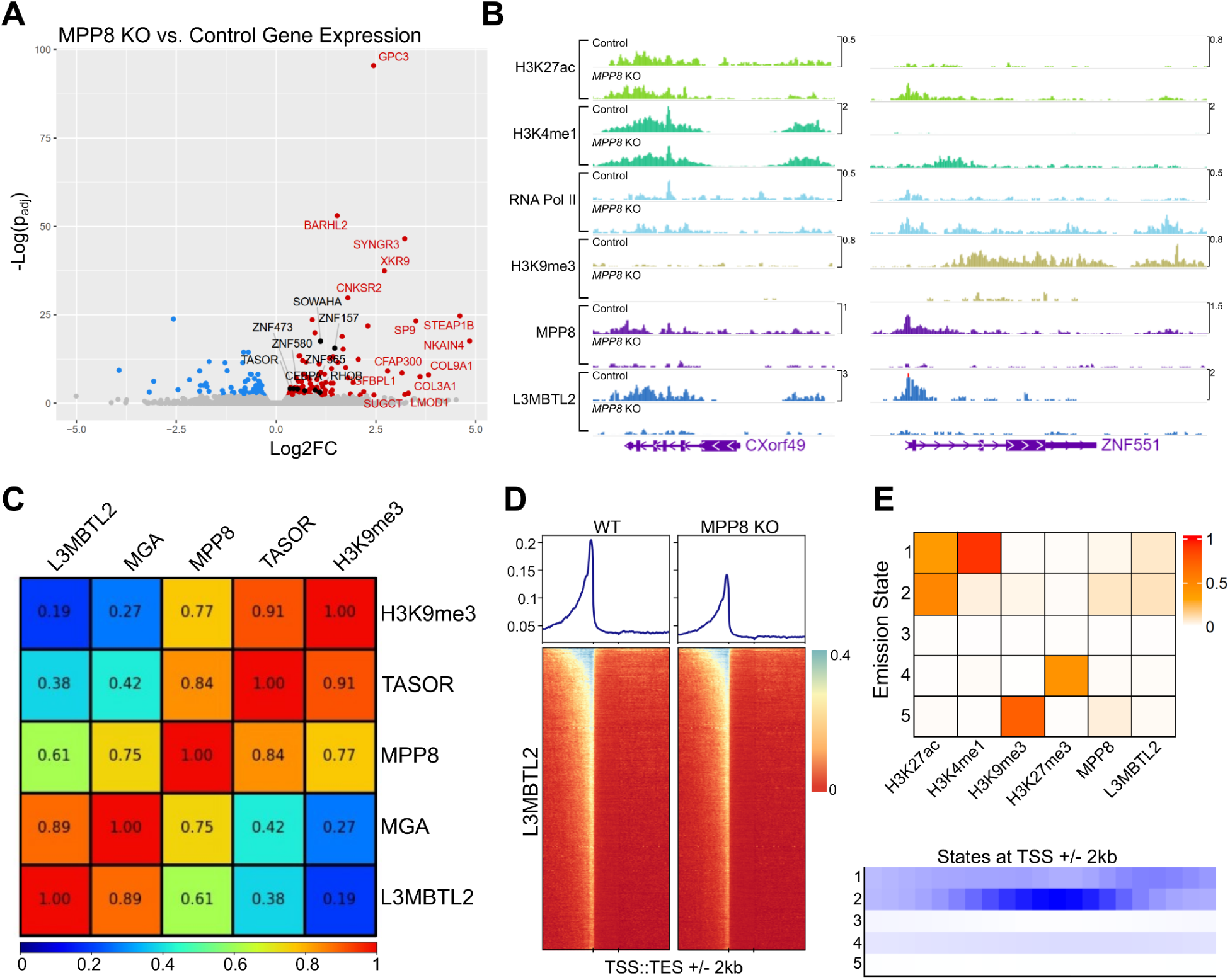
Genome-wide characterization of loci bound by PRC1.6 and HUSH in Jurkat cells. (A) Total RNA sequencing of Jurkat cells harboring *L3MBTL2* knockout versus control Jurkat cells. Colored/labeled points meet significance thresholds (FDR 0.1). (B) Normalized (counts-per-million) CUT&Tag signal targeting named factors at *CXorf49* and *ZNF551* loci in unmodified Jurkat cells (Control) and *MPP8* knockout Jurkat cells (*MPP8* KO). (C) Pearson correlation of counts-per-million-normalized CUT&Tag signal of named factors at annotated human genes (hg38 RefSeq transcription start to end site). (D) L3MBTL2 CUT&Tag coverage map at human genes; transcription start site (TSS) to transcription end site (TES) regions are scaled by length. 2 kilobases upstream and downstream of scaled regions are also depicted. (E) ChromHMM [40] emission states using CUT&Tag from named factors as input (top panel). Coverage of each emission state with respect to transcription start sites (TSSs). For data associated with this figure please see Extended Data Tables 7 and 8 and SRA Bioproject PRJNA869850.

We next examined the chromatin state of endogenous genes upregulated after HUSH knockout in wildtype and *MPP8* knockout Jurkat cells for changes in PRC1.6 binding. To achieve this, we employed CUT&Tag, probing MPP8, L3MBTL2, H3K9me3, and markers of transcriptional activity (H3K4me1, H3K27ac, and RNA Polymerase II). Loci bound by MPP8 in the wildtype background were considered directly HUSH-regulated if, after *MPP8* knockout, they demonstrate decreased H3K9me3 signal and increased transcription marker signal. In a small subset of these genes, we noted that L3MBTL2 signal distribution closely mirrors that of MPP8 and is severely diminished after *MPP8* knockout (Figure 3B and SRA Bioproject PRJNA869850). Together, these transcriptomic and genomic data demonstrate that PRC1.6 localization is HUSH-dependent at a subset of silenced loci, but contributes only modestly to the maintenance of silencing. It is unclear whether, in developing or fully differentiated cells, PRC1.6 is required to establish transcriptional repression at HUSH targets.

### Characteristics of HUSH- and PRC1.6-bound chromatin genome-wide

To understand the frequency with which PRC1.6 and HUSH occupy the same endogenous loci, we examined MPP8, TASOR, L3MBTL2, and MGA CUT&Tag signal genome-wide (Figure 3C and SRA Bioproject PRJNA869850). Within regions between transcription start sites (TSSs) and transcription end sites (TESs), we observed strong correlation between core components of HUSH (cor = 0.84) and PRC1.6 (cor = 0.89). Notably, MPP8 signal was also associated with that from the two PRC1.6 components, MGA (cor = 0.75) and L3MBTL2 (cor = 0.61). H3K9me3 clustered well with HUSH and poorly with PRC1.6, suggesting that the link between the two protein complexes may not be limited to silenced genes. There was a relatively weak association between either complex and H3K27ac, H3K4me1, RNA Polymerase II, and H3K27me3.

We expanded surveyed genic regions by 2 kilobases from TSS/TES to assess the distribution of L3MBTL2 along genes (Figure 3D and SRA Bioproject PRJNA869850). Compared to wild type Jurkat cells, *MPP8* knockout cells exhibit a decrease in L3MBTL2 CUT&Tag signal at aggregate promoters, suggesting that HUSH and PRC1.6 may co-localize to *cis*-regulatory regions rather than gene bodies.

We also sought to stratify the chromatin environments amenable to HUSH and PRC1.6 localization genome-wide. To achieve this, we applied CUT&Tag signal from MPP8, L3MBTL2, and histone marks associated with transcriptional activity or silencing to analysis by ChromHMM [39], limiting output to five emission states (Figure 3E and Extended Data Table 8). We classified states as follows: (1) active enhancers marked by H3K27ac and H3K4me1, (2) promoters marked by H3K27ac signal enriched at TSSs, (3) unmarked or unmappable regions, (4) H3K27me3 constitutive heterochromatin, and (5) H3K9me3 silenced regions. MPP8 signal is most prominent at states 2 and 5, indicating localization to euchromatic promoters and regions silenced by H3K9me3. L3MBTL2 signal was enriched at states 1 (enhancers) and 2 (promoters), without accumulation at state 4 (which is presumably associated with canonical polycomb complexes). We conclude that HUSH and PRC1.6 associate primarily at active promoters rather than at silenced regions.

Previous studies characterizing HUSH complex genomic localization have focused on H3K9me3-silenced regions that undergo transcriptional upregulation after depletion of HUSH core components [4,6,13]. The means by which the HUSH complex is recruited to active *cis* regulatory elements remains largely unexplored. We therefore asked whether loci strongly bound by HUSH are associated with known transcription factor binding sites. Using HOMER [41], we identified known transcription factor motifs enriched in genomic regions with the highest MPP8 CUT&Tag signal (top 10,000 peaks) compared to human promoter sequences. P53, ZNF41, and 4 Homeobox factors are among the highest enriched over background (Supplementary Figure 3 and Extended Data Table 10). We next examined MPP8 peaks strongly associated with L3MBTL2 CUT&Tag signal (n=800 intersecting peaks). In this subset, we found motifs corresponding to 2 of the above Homeobox factors among the most significantly enriched over promoter background (Supplementary Figure 4 and Extended Data Table 11). These results support a model in which HUSH interacts with diverse chromatin modifiers–including polycomb repressive complexes–to regulate gene expression during development [42,43], and exhibits persistent localization to target gene promoters following differentiation, as evidenced in the cell line here.

### The locus-specific interaction between HUSH and PRC1.6 is found in more than one cell line

Finally, we asked whether transcriptional and epigenetic shifts observed in HUSH and PRC1.6 knockout Jurkat cells are reproducible in HEK293T. Jurkat cells were used in the loss-of-function screen that identified the PRC1.6 components L3MBTL2 and MGA. HEK293T cells were used in the Provirus Proximal Proteomics screen that identified L3MBTL2 near the HUSH-silenced provirus. In *MPP8, TASOR, PPHLN1,* and *L3MBTL2* mutants, we observed significant (p_adj_ < 0.05, log2FC > 1) upregulation of 90, 69, 68, and 62 genes, respectively. Among upregulated genes, *RARRES2P2*–a pseudogene–was upregulated in all knockouts (Figure 4A and Extended Data Table 9). HUSH-mediated silencing of pseudogenes and retrogenes is observed across cell types, as discussed in [10]. We also compared *L3MBTL2* knockout transcriptomes from HEK293T and Jurkat cells directly and found *LHX6*, *SYNGR3*, *ZNF157*, *GPC3*, *XKR9* are upregulated in both lines (Figure 4B and Extended Data Table 9). Consistent with previous results [6], *MPP8* knockout resulted in widespread transcriptional upregulation within the chromosome 19 ZNF gene cluster (Figure 4C and Extended Data Table 9). MPP8 and L3MBTL2 genomic localization within this region revealed overlap between the factors primarily at promoters, as we found in Jurkat cells. Moreover, L3MBTL2 localization to MPP8-bound chromatin appears to be highly dependent on the latter factor. These results demonstrate that the locus-specific interaction between PRC1.6 and the HUSH complex, at HUSH-repressed genes, is reproducible across cell lines.

**Figure 4:**
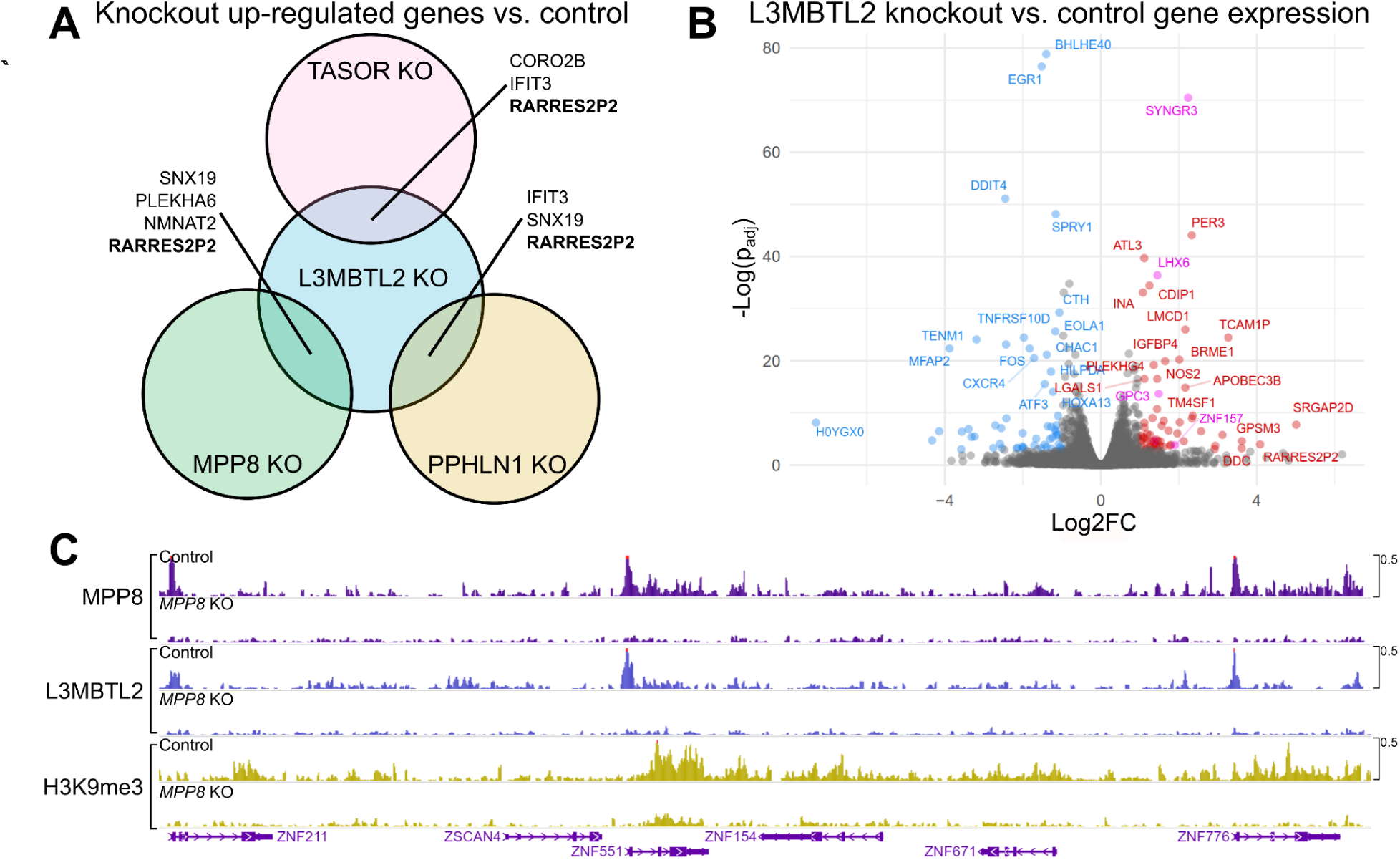
Transcriptome-wide and genome-level effects of HUSH and L3MBTL2 depletion in HEK293T cells. (A) Venn diagram depicting genes that are upregulated vs control (p_adj_<0.05, log2FC>1) in *TASOR*, *MPP8*, *PPHLN1*, and *L3MBTL2* knock out HEK293T cells. (B) Volcano plot showing differential gene expression (p_adj_<0.05, log2FC>1) in *L3MBTL2* knockout cells compared to control knockouts. Colored points depict genes meeting the significance threshold. Magenta points indicate genes that are also upregulated in Jurkat *L3MBTL2* knockouts. (C) *MPP8*, *L3MBTL2*, and H3K9me3 CUT&Tag in control knockouts and *MPP8* knockouts. Genomic region depicted is the chromosome 19 *ZNF* gene cluster. For data associated with this figure please see Extended Data Table 9 and SRA Bioproject PRJNA869850.

## DISCUSSION

In this study, we developed a method of spatially-restricted tagging of proteins at a newly-integrated provirus (Provirus Proximal Proteomics). The exact screen used here was designed such that the provirus is transcriptionally silenced by the HUSH complex. Proteins associated with the provirus were identified both in the presence and in the absence of HUSH. To our knowledge, this is the first assay using retroviral DNA as bait to identify interacting proteins involved in transcriptional repression. The results illuminate a complex interplay between proteins that support and inhibit viral transcription and replication. Even in the presence of efficient HUSH silencing, a host of transcription proteins and RNA processing proteins were recruited to the provirus. By comparing enriched proteins with forward genetic screens for lentiviral transgene silencing factors, we found that transgene repression is jointly dependent on the non-canonical polycomb repressive complex 1.6 (PRC1.6)–a complex enriched by C-BERST–as well as HUSH. Genome-wide mapping of HUSH, PRC1.6, and key histone post-translational modifications revealed frequent co-localization at endogenous promoters across the genome without evidence of repressive activity from either complex. Overall, our work builds a more nuanced picture of HUSH complex protein interactions and function, and raises further questions about the mechanisms orchestrating durable silencing of exogenous genetic elements.

### HUSH establishes a complex chromatin environment at proviruses

Proximity labeling at a HUSH-silenced provirus revealed numerous transcription and mRNA processing proteins, suggesting that even while repressed, these exogenous elements remain targeted by transcriptional machinery. This is perhaps not surprising since recruitment of the HUSH complex to mediate transcriptional silencing is dependent upon transcription [10], and, as demonstrated here and elsewhere [3,6,13,44], HUSH complex proteins associate with promoters across the genome (Figure 3E). In the context of robust reporter silencing in HUSH^+^ cells (Supplementary Figure 1), we also found proteins that facilitate DNA methylation, DNA repair, histone deacetylation, and recruitment of heterochromatin-associated proteins. The dynamic competition between activating and repressing complexes likely underpins the cell type- and state-specific responses observed at HUSH-targeted loci. Further studies should explore how various factors may change in relative abundance according to cellular context.

### PRC1.6 is associated with HUSH complex-silenced proviruses

The finding that L3MBTL2, a PRC1.6 component, is enriched at silenced proviruses was unexpected given that HUSH predominantly mediates transcriptional silencing via an H3K9me3-driven repression pathway [3], while components of polycomb complexes generally catalyze mono-, di-, and tri-methylation of H3K27 [45–47]. Nevertheless, the findings here are supported by a large-scale proteomics study in which networks of histone post-translational modifications and their associated protein complexes were deconvolved [48]. Specifically, these researchers found that H3K9me3 is associated with protein components of the HUSH, PRC1.6, and ORC complexes. Interestingly, components of the latter two complexes - L3MBTL2 and ORC5, respectively - were enriched at the provirus in our Provirus Proximal Proteomics screen. The variant histone H2AZ.2 and VPS72, the chaperone that promotes H2AZ.2 deposition on chromatin [49,50], were also both associated with the provirus in the HUSH^+^ cells. The latter two hits are particularly interesting since H2AZ.2 is a component of nucleosomes decorated with H2K119ub, a histone post-translational modification that is deposited by PRC1.6 [51].

Our forward genetic screen provided orthologous evidence for the relevance of PRC1.6 in the repression of HUSH complex-sensitive proviruses. Interestingly, as we showed previously for HUSH complex-mediated silencing [14], the identity of the promoter influences the sensitivity of an otherwise isogenic reporter gene to silencing by PRC1.6 (Figure 2D). Perhaps this effect is due to differences in the level of transcription, a determinant of sensitivity to silencing by the HUSH complex [10]. One possibility is that PRC1.6 is recruited to HUSH-silenced loci independent of H3K9 modification state and plays a subtle role in repressing transcription, shaping chromatin architecture, or both. It is also possible that the silencing function of PRC1.6 extends primarily to the establishment of transcriptional repression during differentiation or genomic integration of new foreign DNA elements, rather than to maintenance of silencing at endogenous loci.

### PRC1.6 and HUSH co-localize at endogenous chromatin targets

While L3MBTL2 knockdown had no observable effect on HUSH-repressed endogenous gene expression in Jurkat cells, our genomic binding analysis using CUT&Tag indicated a striking co-occurrence of HUSH and PRC1.6 components at a subset of genomic loci. The loss of L3MBTL2 signal in *MPP8* knockout cells suggests that the localization of PRC1.6 at some repressed genes is dependent on the HUSH complex. Furthermore, ChromHMM analysis revealed that the preferred genomic context for shared HUSH and PRC1.6 targets is euchromatic promoters. Despite prolific promoter binding genome-wide, it is intriguing that transcription of most downstream genes remains unchanged after depleting core components of either complex. At regions bound strongly by the HUSH complex alone, or by both the HUSH complex and PRC1.6, there was notable enrichment of transcription factor binding motifs corresponding to developmental (Homeobox) factors. This suggests that the biological roles of the HUSH complex and PRC1.6 are likely more obvious during development and merit investigation within experimental systems for development.

There are several plausible mechanisms by which HUSH and PRC1.6 may be recruited to the same genomic regions: (1) HUSH and PRC1.6 transcriptionally repress endogenous genes during the developmental program in SETDB1-dependent fashion. The colocalization of these two protein complexes to promoters in differentiated cells may therefore be a remnant of the developmental program (Figure 5A) [1,52]. (2) We and others have demonstrated that the degree of transcriptional silencing by HUSH is promoter-specific [3,8,14,21], and extend this observation to PRC1.6. Intrinsic properties of select exogenous and endogenous promoters such as viral origin, paused RNA Polymerase II occupancy, or transcription factor binding likely contribute to localization or activity of PRC1.6 and the HUSH complex. In the endogenous context, these complexes may safeguard active promoters from perturbation by transposon and retrovirus integration (Figure 5B). (3) L3MBTL2 binds directly to H3K9 via its MBT domain, perhaps after the HUSH complex triggers H3K9 methylation via SETDB1 or G9a (Figure 5C) [3,53]. (4) HUSH complex protein localization may act directly, or indirectly through DNA damage, to recruit DNA repair machinery such as MDC1, ERCC1, FANCI, and SMC5, each of which was found by Provirus Proximity Proteomics at the provirus in HUSH^+^ cells. MDC1 was also found to cooperate with HUSH as part of rDNA repair, and depends on L3MBTL2 for recruitment of accessory repair factors (Figure 5D) [33,54].

**Figure 5:**
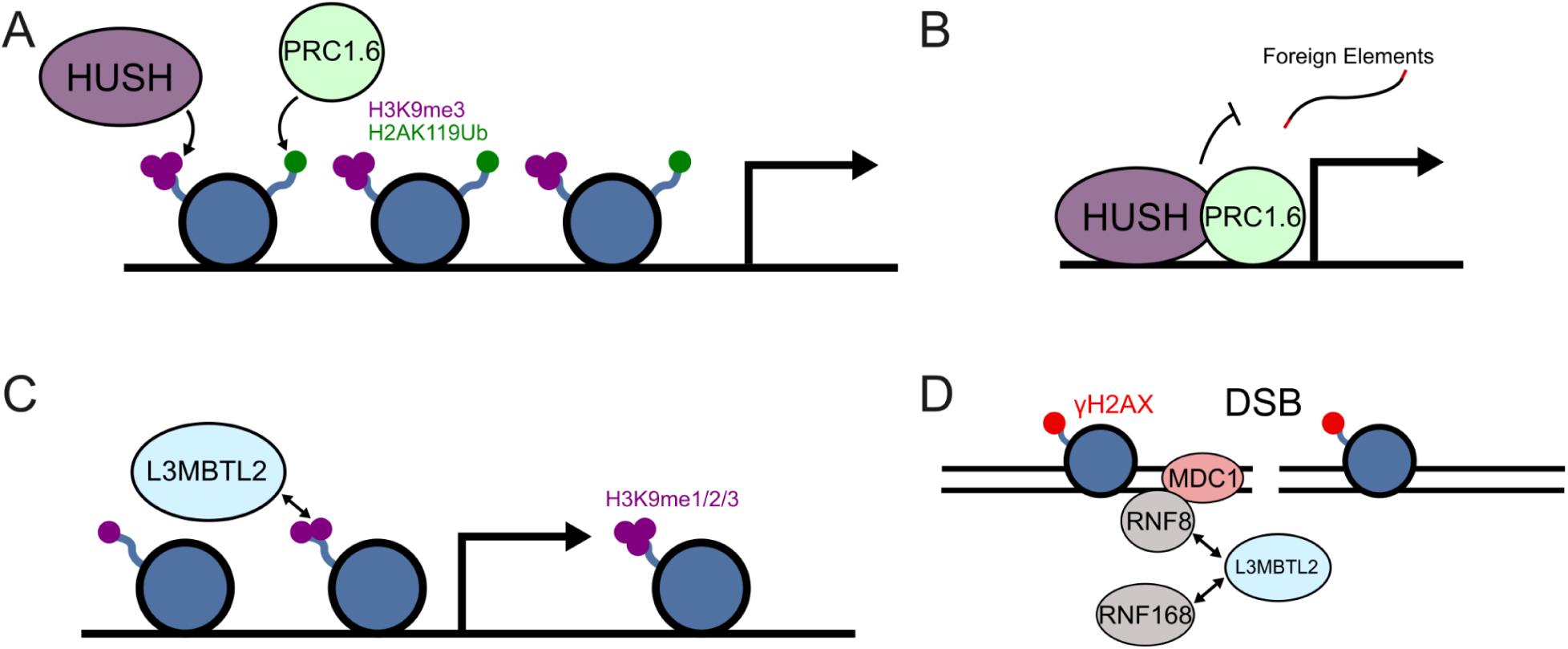
Possible mechanisms underlying co-localization of HUSH and PRC1.6 complexes. (A) The HUSH and PRC1.6 complexes transcriptionally repress developmental genes and overlap at select loci. (B) Both complexes reside at endogenous promoters to safeguard against perturbation by retroviruses and retrotransposons. (C) L3MBTL2 may be recruited to HUSH loci through recognition of H3K9me3 marks previously established by the HUSH complex. (D) DNA damage may independently recruit the HUSH and PRC1.6 complexes to overlapping loci (adapted from [54]).

## MATERIALS AND METHODS

### Plasmids

All plasmids used in this study are listed with their corresponding Addgene accession numbers in Extended Data Table 12. The plasmids themselves, along with their complete nucleotide sequences, are available at https://www.addgene.org/Jeremy_Luban/.

### Cell culture

HEK293T cells were obtained from ATCC (CRL-3216) and cultured in DMEM + 10% FBS + Penicillin-Streptomycin (100U/mL). Jurkat cells were obtained from ATCC (TIB-152) and cultured in RPMI + 10% Penicillin-Streptomycin (100U/mL).

### Provirus Proximity Proteomics

dCas9–APEX2 biotinylation at genomic elements by restricted spatial tagging (C-BERST) was performed as described [55] and modified for the identification of proteins associated with provirus. HEK293T cells were first transduced with a lentivector that had packaged RNA from pEJS578_DD-dSpyCas9-mCherry-APEX2. Then sgRNAs were delivered using a Sleeping Beauty vector by co-transfecting pCMV(CAT)T7-SB100 with plasmids expressing sgRNAs from the U6 promoter: pSB_EJS614_pTetR-P2A-BFPnls/sgLTR encoded an sgRNA targeting the provirus LTR and pSB_EJS614_pTetR-P2A-BFPnls/sgNS encoded a non-specific sgRNA. sgLTR was screened for off-target sites using Cas-OFFinder [56]. A population of high BFP-, low mCherry-expressing cells were selected. Cells were then transduced with shRNA vectors to knockdown either *MPP8* or Luciferase. We used three 15 cm dishes of HEK293T cells grown to confluence (20 x 10^6^ cells per dish) per replicate per condition ([sgNS, sgLTR] x [wildtype, *MPP8* shRNA knockdown]) with three replicates for each condition. All plasmids used here and sequences used for specific sgRNAs and shRNAs are described in Extended Data Tables 12 and 13.

As described in [55], dCas9-mCherry-APEX2 expression was induced after treatment with doxycycline and Shield1 ligand (Takara 632189) for 24 hours and 16 hours, respectively. Cells were then treated with biotin-phenol (Adipogen, cdx-b0270). APEX2-mediated proximity labeling was then induced by the addition of H_2_O_2_ and quenched with a sodium ascorbate solution, followed by PBS washes. After nuclear isolation, sonication, and isolation of biotinylated proteins on streptavidin dynabeads, proteins were eluted using SDS loading buffer and submitted for mass spectrometry. Key reagents used here are described in Supplementary Table 14 and in reference [57]. iBAQ scores for each condition were calculated based on enriched peptides in three replicates per condition. We excluded peptides that were found in only one replicate. To calculate Log2 fold-change of iBAQ scores and rank-adjusted p-values (p_adj_) of sgLTR/sgNS enrichment for both wildtype and knockdown, we used the DESeq package in R [58] with scale set to 1 (avoiding adjustment for transcript length).

### Genome-wide CRISPR-Cas9 Screen

The all-in-one (Cas9, puromycin selection cassette, and sgRNA library) Toronto KnockOut (TKO) CRISPR library Version 3 (LentiCRISPRv2 backbone with Pooled sgRNA Library), was a gift from Jason Moffat. To prepare sgRNA library-containing lentivirus, HEK293T cells were transfected with library plasmid DNA, psPAX2, and pMD2.G. To generate pools of knockout Jurkats, 40 x 10^6^ cells were transduced with lentiviruses containing a library at an MOI of 0.3, followed by selection in puromycin, either 1 ug/mL or 10 ug/mL, for two weeks. The brightest GFP cells (top 5%) were sorted by FACS and recovered. Cells were expanded and sorted a second and a third time to enrich for high GFP-expressing cells. Genomic DNA was extracted from the sorted cells and sgRNAs were sequenced on a NextSeq550 in single-end mode and compared to the sgRNA pool of unsorted cells passaged concurrently with the sorted experimental group. The sgRNA counts and abundance were analyzed using MAGeCK [37].

### Generation of knockout cell lines and confirmation of screen hits

Pooled Jurkat knockouts were generated by Invitrogen NEON mediated electroporation of 3xNLS-SpyCas9 protein complexed with the indicated synthetic sgRNAs (Cas9-gRNA RNP) ordered from IDT (Alt-R based guide). 3xNLS-SpyCas9 protein was bacterially expressed and purified as previously described [59]. 80 nmol of 3xNLS-SpyCas9 was incubated with 100 nmol of guide in 6 uL of R buffer. After 15 minutes 5 x 10^5^ Jurkat cells in 6 uL of R buffer were mixed with the preformed Cas9-gRNA RNP complexes and electroporated using the NEON system using suggested settings for Jurkat cells. After electroporation, cells were recovered and grown and tested for knockout efficiency by western blot analysis of proteins encoded by the indicated genes.

To confirm the effect of *L3MBTL2* on provirus silencing, Jurkat cells were transduced with lentiviral vectors bearing pEF1a-Zim3-dCas9-P2A-EGFP or pSFFV-Zim3-dCas9-P2A-EGFP, two previously described reporter plasmids [38] that were a gift from Marco Jost and Jonathan Weissman (Addgene plasmids #188778 and #188898, respectively). Each population was treated with shRNA vectors targeting *MPP8* (left panels) or *L3MBTL2* (right panels) in blue, vs the non-targeting shRNA control in red. (E) Normalized CUT&Tag signal (counts-per-million) for MPP8, L3MBTL2, and H3K9me3 at the pscALP-PLXIN-gagGFP-WPRE provirus in HEK293T cells.

### Cleavage Under Targets and Tagmentation (CUT&TAG)

100,000 viable nuclei were isolated from freshly cultured Jurkat cells used for CUT&Tag (Cleavage Under Targets and Tagmentation) [60]. Following incubation with primary antibodies, samples were incubated with a secondary anti-rabbit IgG antibody (Novus NBP172763). Libraries produced by tagmentation and barcoding were sequenced on a NextSeq550 in paired-end mode (38/37 cycles). Reads were aligned to GRCh38 with BWA-MEM [61]. BigWigs were normalized by counts-per-million (CPM) using Deeptools2 BamCoverage with bin size set to 10 [62].

### Transcriptome analysis

Total RNA sequencing, paired-end (150/150 cycle) with rRNA depletion, was carried out by Azenta. FASTQ files served as input for transcript quantification by kallisto [63] (-b 100) aligned against gencode v40 complete transcriptome reference. Differential gene expression analysis was conducted using sleuth with p-value adjustment as described [64].

### Data Accessibility

Raw RNA-Seq and CUT&Tag files are available under SRA Bioproject PRJNA869850. Mass spectrometry output (tab-separated format) is available in the Extended Data file with this manuscript.

## ACKNOWLEDGEMENTS

We thank members of the Luban and Sontheimer Labs for helpful comments and support, and Stephen Goff, Zsuzsanna Izsvak, Marco Jost, Jason Moffat, Didier Trono, and Jonathan Weissman for plasmids. This work was funded by NIH grants R37AI147868 and U54AI170856 to J.L., UM1 HG011536 to E.J.S., and R01HL150669 to S.A.W.

## AUTHOR CONTRIBUTIONS

TCR, LY, EJS and JL designed the experiments described in this manuscript. TCR and LY executed “C-BERST” proteomics experiments. LY performed CRISPR knockout screens. LY, NK, OL, EJ, SAM, and SAW generated knockout cell lines. LY, NK, OL, and EJ performed virus transduction and silencing experiments. TCR, LY, and EJ performed RNA-sequencing and CUT&Tag experiments. TCR conducted analysis of proteomics and sequencing data. SAM and SAW purified and QCed the SpyCas9 protein

---

**Supplementary Figure 1:**
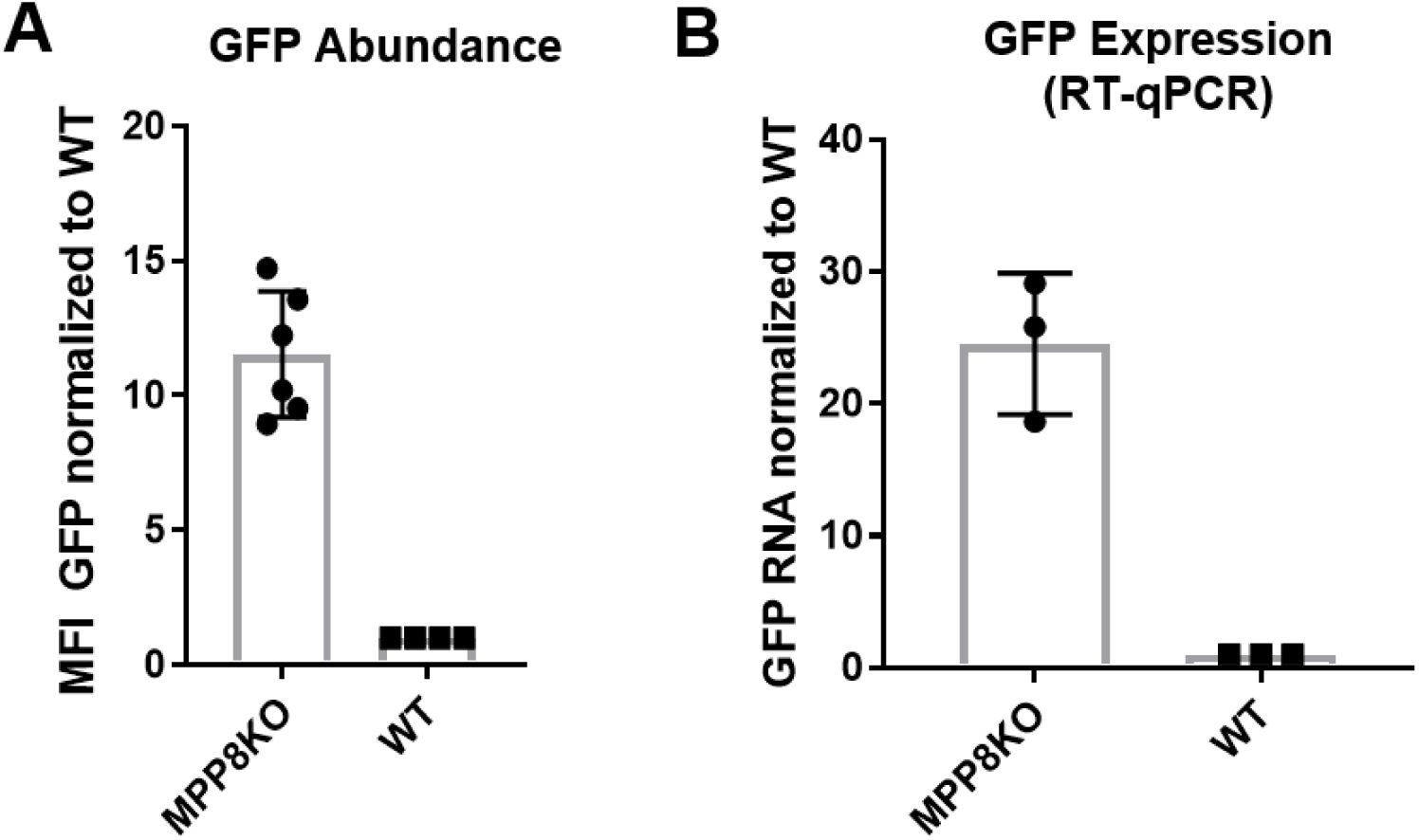
Transcriptional repression of pscALP-PLXIN-gagGFP-WPRE by the HUSH complex. (A) GFP mean fluorescence intensity (MFI) of control (WT) or MPP8 knockout (MPP8KO) HEK293T cells transduced with pscALP-PLXIN-gagGFP-WPRE. (B) RT-qPCR enrichment of pscALP-PLXIN-gagGFP-WPRE RNA in HEK293T control (WT) or MPP8 knockout (MPP8KO) HEK293T cells transduced with pscALP-PLXIN-gagGFP-WPRE.

**Supplementary Figure 2:**
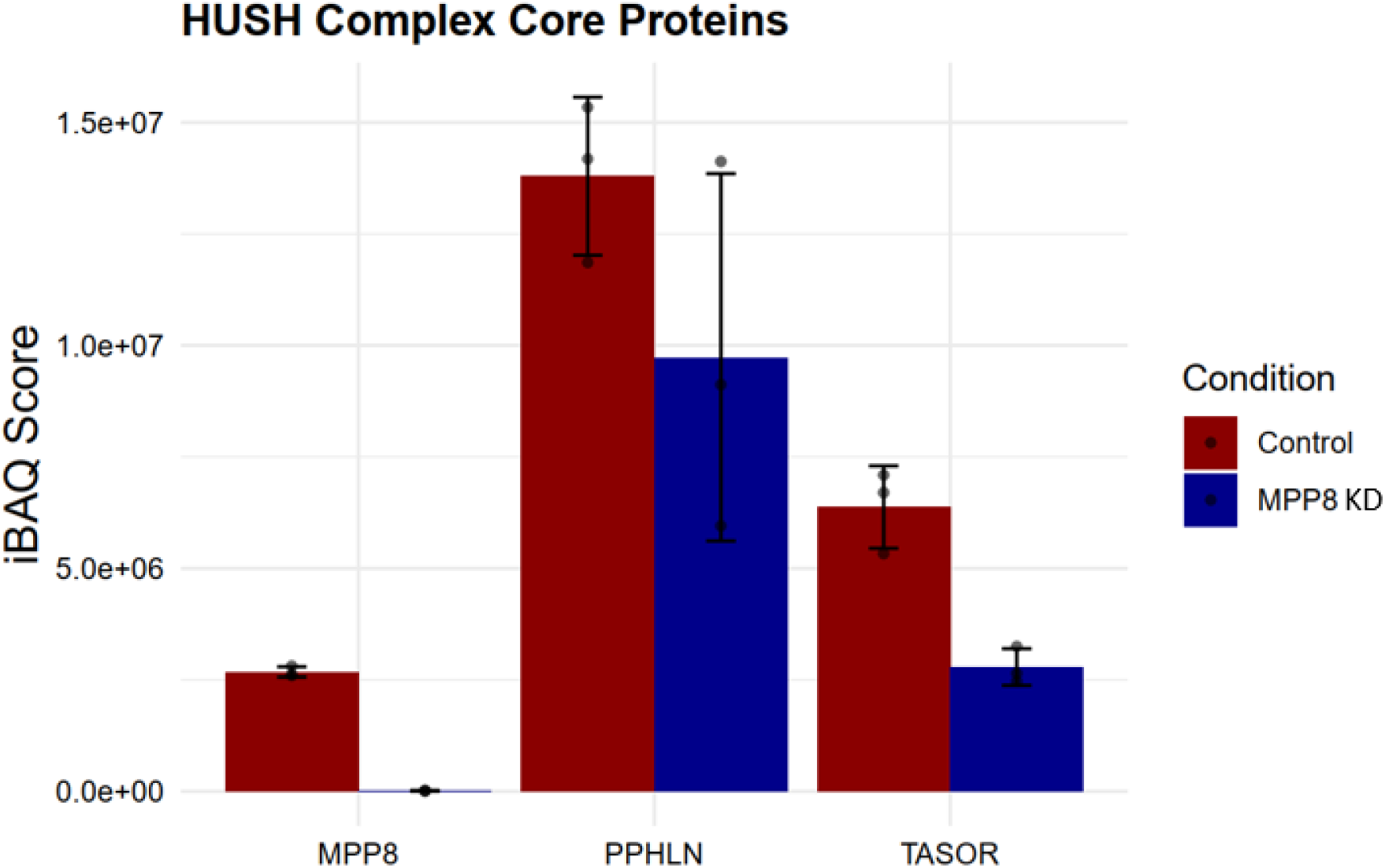
HUSH core proteins detected by Provirus Proximity Proteomics. HEK293T cells were transduced with dCas9-APEX2 and expressed sgRNA targeting the lentiviral provirus (sgLTR). Control cells were transduced with an shRNA targeting luciferase. shMPP8 cells are transduced with an shRNA targeting MPP8 (core HUSH component). iBAQ values across 3 biological replicates and depicted for control and *MPP8* knockdown conditions. For data associated with this figure please see Extended Data Table 1.

**Supplementary Figure 3:**
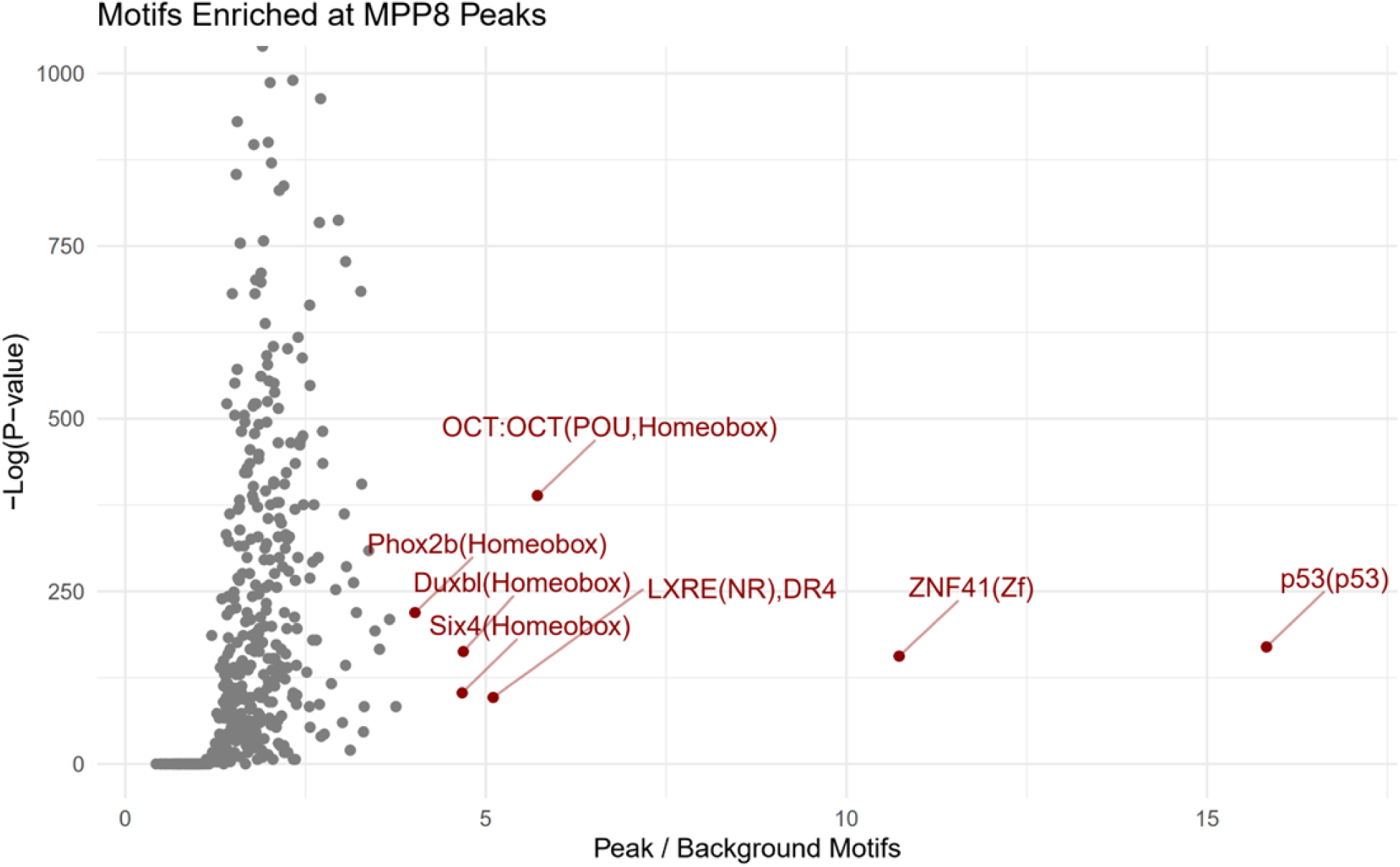
Motif enrichment at MPP8 peaks. De novo motif discovery applied to DNA sequences of top 10,000 MPP8 peaks (by CUT&Tag signal), subtracting “simple repeat” regions (RepeatMasker) likely to result from spurious mapping. 24.6% of peaks overlap with annotated transcription start sites. Genomic sequences from 10,000 randomly selected “promoter-like” cis regulatory elements (ENCODE) were used as background signal. Points represent individual motifs. X axis values represent the proportion of motif occurrences in the test (MPP8) regions over those found in the background regions. Y axis values are the –Log(p-value) calculated by Homer. Red points depict motifs x > 4 and p < 0.001. For data associated with this figure please see Extended Data Table 10 and SRA Bioproject PRJNA869850.

**Supplementary Figure 4:**
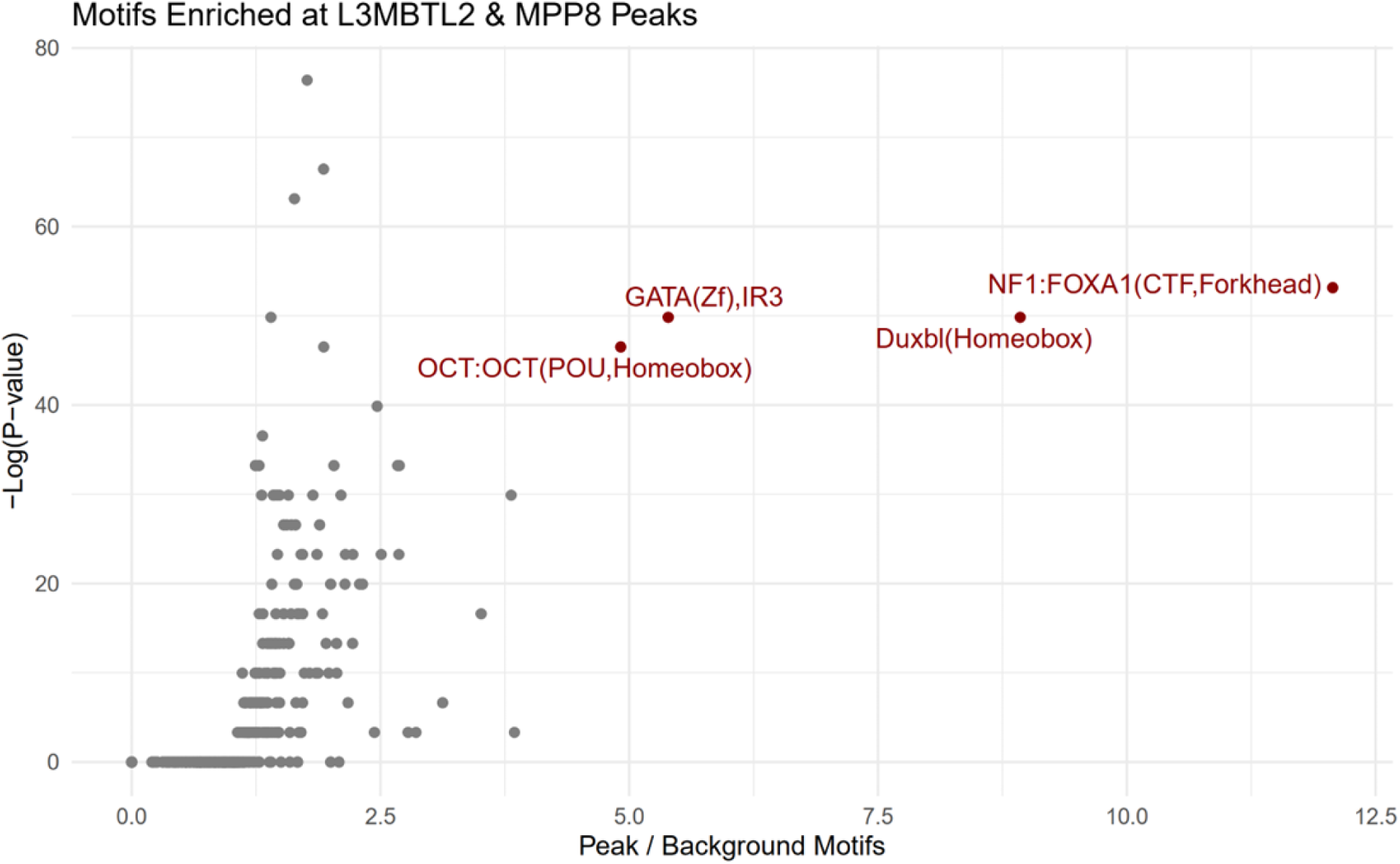
Motif enrichment at L3MBTL2 and MPP8 peaks. De novo motif discovery applied to DNA sequences of intersecting regions from top 10,000 MPP8 peaks and top 10,000 L3MBTL2 peaks (by CUT&Tag signal), subtracting “simple repeat” regions (RepeatMasker) likely to result from spurious mapping. 800 overlapping peaks from both factors intersect. 56.75% of peaks overlap with annotated transcription start sites. Genomic sequences from 10,000 randomly selected “promoter-like” cis regulatory elements (ENCODE) were used as background signal. Points represent individual motifs. X axis values represent the proportion of motif occurrences in the test regions over those found in the background regions. Y axis values are the –Log(p-value) calculated by Homer. Red points depict motifs x > 4 and p < 0.001. For data associated with this figure please see Extended Data Table 11 and SRA Bioproject PRJNA869850.

